# Skin microbiota dynamics of dairy cows affected by digital dermatitis

**DOI:** 10.1101/2019.12.20.882787

**Authors:** Juan Manuel Ariza, Dörte Döpfer, Kenny Oberle, Kelly Anklam, Sophie Labrut, Nathalie Bareille, Anne Relun, Raphaël Guatteo

## Abstract

Bovine digital dermatitis (DD), is a highly prevalent disease among dairy cattle characterized by ulcerative and painful lesions. While multiple management factors are involved in the disease, its precise etiology remains uncertain and the effectiveness of current control strategies remains highly variable. The major role of *Treponema* spp. in the development of the disease is consistently recognized. Nevertheless, it remains unclear how other bacterial communities are relevant to the onset and progression of the disease, and how the skin microbiota is affected by the environment during the course of the disease. The objective of this study was to describe the dynamics of microbiota recovered from DD affected feet under field conditions. This study described the diversity, structure, and composition of DD lesion microbiota over 45 days according to different clinical and management factors. The results of this investigation confirmed the existence of a specific skin microbiota associated with DD lesions, dominated by *Treponema* spp. and very different from the microbiota of healthy skin. Interestingly, the diversity and structure of the microbiota in DD lesions did not vary with the footbath disinfectant or the individual topical antibiotic treatments used. In addition, microbiotas from proliferative lesions evidenced a different structure and diversity in comparison to non-proliferative lesions. Our results confirm the major role of *Treponema* spp. And highlight the potential role of *Mycoplasmopsis* spp. in the DD lesion onset. Further studies are needed to confirm whether the clinical course of DD lesions is driven by a particular microbiota and how that microbiota may induce disease.

**Highlights:** Multiple bacteria have been identified in DD lesions. However, many of these microorganisms are inhabitants of the foot skin and the farm environment. For the first time, the microbiota of DD lesions was monitored for 45 days under field conditions to describe its evolution over time. The results of this investigation highlighted a particular microbiota dominated by *Treponema* spp. present on the skin of DD affected animals and highly different from those microbiotas of healthy skin. The microbiota of DD lesions evolved over the study period and differential bacteria were identified. Further studies are warranted to determine the role of the bacteria composing these microbiotas on lesion onset and outcome.

## Introduction

The increased prevalence of lameness and its impact on the welfare and productivity of animals reflects an important concern facing modern dairy farming. One of the main causes of infectious lameness is bovine digital dermatitis (DD)(1), a multifactorial and polymicrobial disease spread across the world and characterized by ulcerative and painful lesions which might persist as a chronic condition. The DD lesions progress in dynamic ways conditioned by multiple environmental factors that influence the spread of associated-pathogens and the integrity of the foot skin. Currently, the control strategies implemented in dairy farms such as footbaths or individual treatments, have demonstrated variable effectiveness in the healing and/or prevention of DD lesions(2, 3).

Although the precise etiology of DD remains unclear, the previous maceration of the foot skin and the presence of *Treponema* spp. have been consistently identified as the major etiological components of DD(4–7). However, multiple other bacteria have also been associated with DD infection in different studies, such as *Mycoplasma* spp.(8, 9); *Fusobacterium necrophorum*(10); some Bacteroides, specifically *Porphyromonas levii*(11, 12); different Proteobacteria, such as *Campylobacter* spp. and *Dichelobacter nodosus*(13–15), and finally a broad range of other Firmicutes(16). Nevertheless, the large part of these DD-associated pathogens (including the *Treponema* spp.) are inhabitants of the normal foot skin, the rumen, and the gastrointestinal tract of ruminants, and thereby, of the farm environment(17–19). Together, all these findings have reinforced the concept of DD as a polymicrobial disorder suggesting that a particular microbiota may drive the lesion environment and affect its outcome.

The development of collective and individual treatment strategies has been focused on the incriminate DD associated-pathogens, mainly against *Treponema* spp. Nevertheless, these treatments may influence the structure of the microbial communities that constitute the foot skin barrier. The disruption of the normal skin microbiota can lead to dysbiosis and consequently induce colonization of DD-associated pathogens and the consequent apparition of DD lesions. Hypothetically, a successful control strategy should regulate microbiota for preventing or decreasing the proliferation of the DD-associated pathogens.

Next-generation sequencing (NGS) had enlarged the perspectives to explore deeply the dynamics of the entire skin microbiota, beyond the restrictions imposed by the culture of pathogens or the pre-selection of targeted species. Through the sequencing analysis of the 16S ribosomal RNA (rRNA), this investigation explored the dynamics of DD microbiotas over 45 days using skin biopsies collected from 10 cows of 5 different dairy farms, and submitted to recognized treatment strategies (individual topical treatment and/or footbath disinfectant)

## Results

A clinical and microbial description of the lesions studied is presented in Table 1 according to the sample time. During the 45 days of follow-up, all 10 initial DD lesions studied remained among the clinical stages M2, M4 or M4.1. None of the DD lesions evidenced a transition to a healing or healthy status (M3-M0). Regarding the lesion type recorded, only two additional proliferative lesions were recorded during the study period. During the study period, 4 feet were treated with topic antibiotic treatment at days 17 (n=2), 35 and 39, respectively. According to the histologic evaluation of epidermal damage, the 10 samples of healthy skin were considered at score 1 and most of the DD lesions were considered at score 3 (n=32), as expected (Table 1). Considering the epidermal damage, while the highest score (EDS=3) was systematically associated with bacterial invasion of spirochaetal origin (Figure 1), lesion with medium score (EDS=2) where less frequently associated with bacterial invasion (BIS=1 (n=3), BIS=2 (n=3), and BIS=3 (n=1)) (Table 1). Finally, lesions with low score (EDS=1) were never associated with bacterial invasion.

**Table 1.**
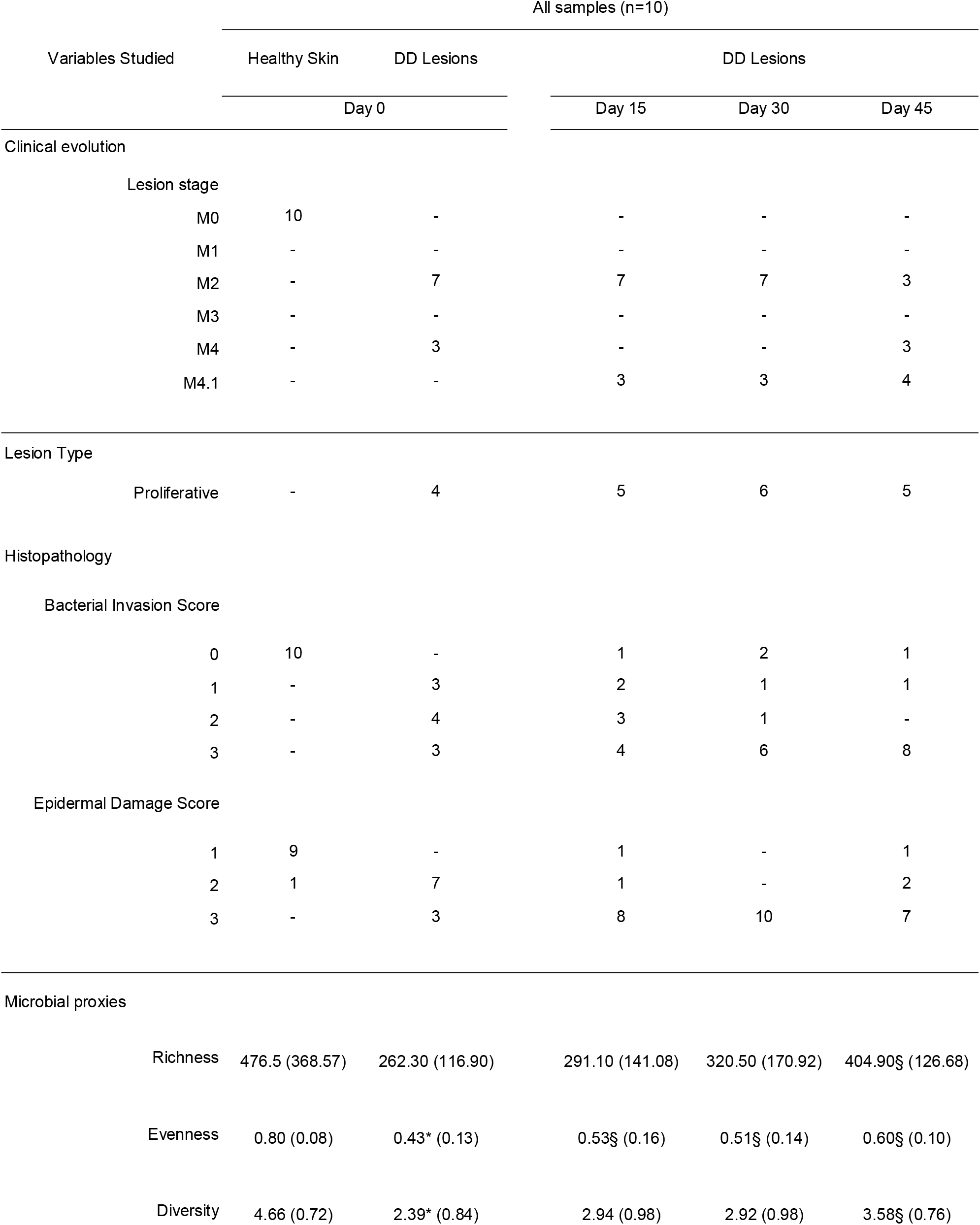
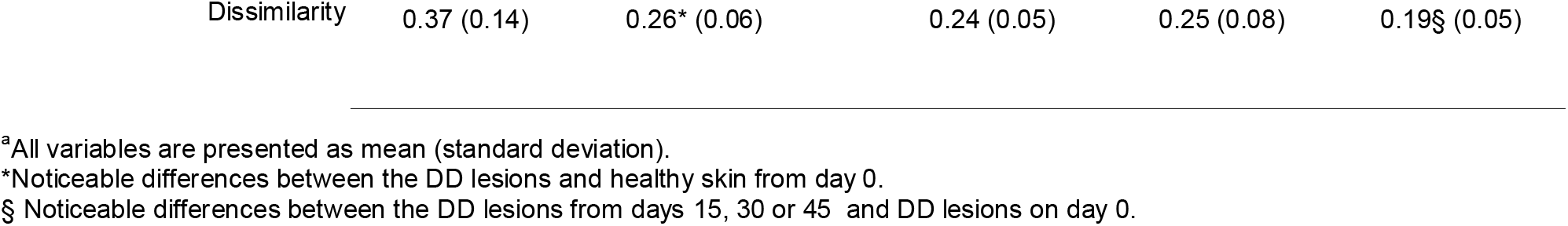
Clinical variables and microbial proxies studied in 10 dairy cows with bovine digital dermatitis lesions followed over 45 days.

**Figure 1.**
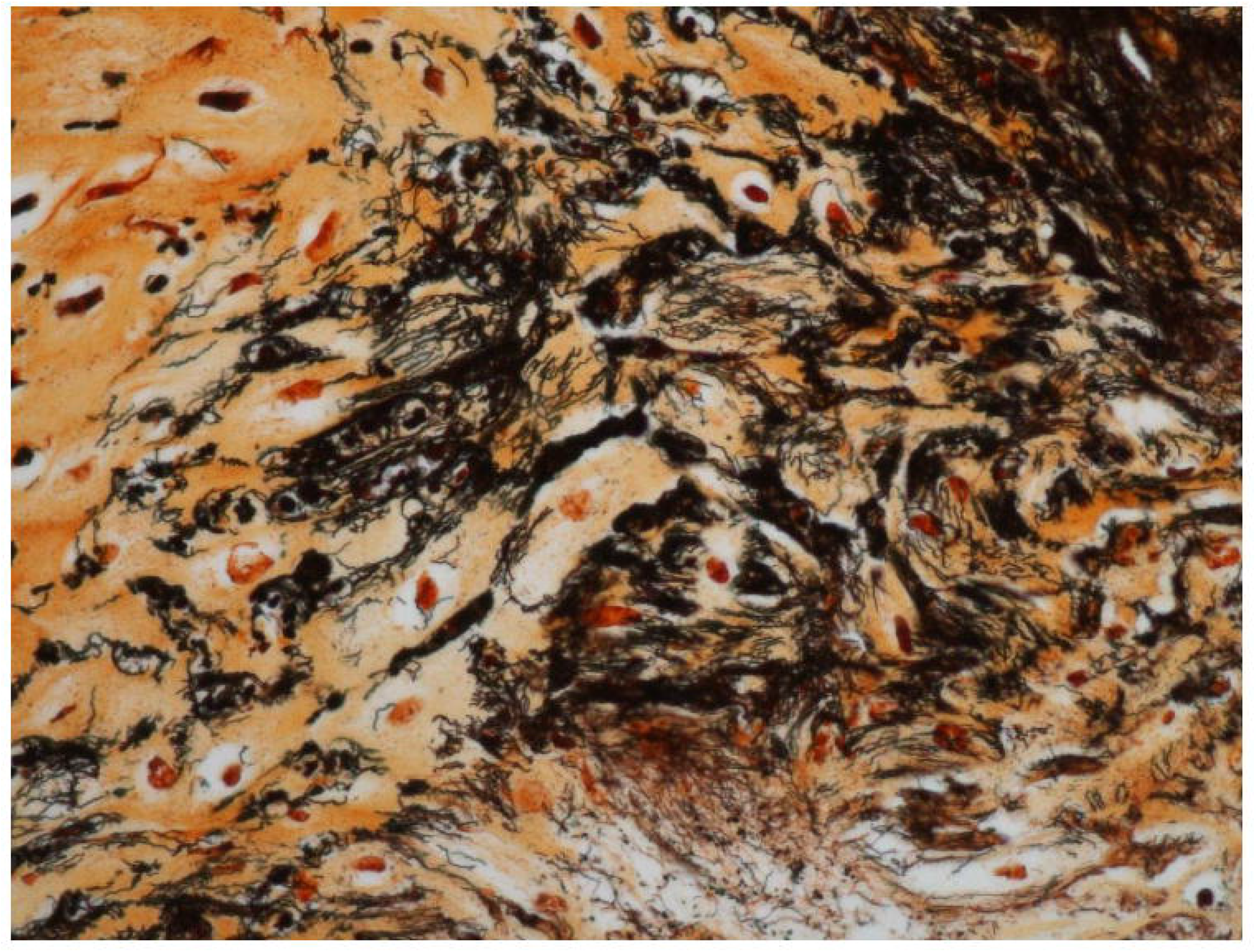
Skin biopsy from a dairy cow with an active bovine digital dermatitis lesion (M2). Colonizing bacteria within surface lesions and defined intra-lesional spirochetes within inflammatory loci are easily evidenced with the Warthin Starry silver staining.

Regarding the raw reads processing, 1571 non-chimeric unique amplicon sequence variants (ASV) were identified. After removing low prevalence ASV occurring in less than two samples, 1398 ASV were conserved. Finally, the phylogenetic tree-based agglomeration conducted to the identification of 139 unique ASV including 8 uniquely named taxa on the phylum level and 24 on the genus level (Figure S1). The set of microbial proxies studied according to every sampling time are presented in Table 1. For the within day of sample comparisons, no differences were found in any of the microbial proxies studied at the start of the study (Healthy skin at day 0, DD lesion at day 0), or during the study (day 15, day 30, and day 45). Otherwise, microbiota from healthy skin showed significantly increased evenness (p = 0.001), diversity (p = 0.001), and dissimilarity (p = 0.027) compared to microbiota from DD lesions sampled at day 0. The evenness of DD lesion microbiota on days 15 and 30 was significantly increased compared with that on day 0. Furthermore, compared to microbiota of DD lesions at day 0, microbiotas collected at day 45 have significantly reduced dissimilarity and increased richness, evenness and diversity.

According to the clinical variables explored and the histological evaluation, no differences in the richness were detected for any of the comparisons (Table 2). In this study, we were unable to detect any significant impact on the evenness, diversity or dissimilarity of the microbiota according to the usage or not of the individual topical treatment and/or the footbath disinfectant. Otherwise, proliferative lesions presented decreased dissimilarity and increased evenness and diversity in comparison to non-proliferative lesions. The DD lesions considered at M2, M4 or M4.1 stages did not differ between them in any of the microbial proxies studied. As expected, the microbiota from healthy skin showed significantly increased evenness, diversity, and dissimilarity compared to any other lesion. Similarly, microbiota from samples considered at the lowest BIS and EDS scores, had noticeably higher evenness, diversity, and dissimilarity (Table 2).

**Table 2.**
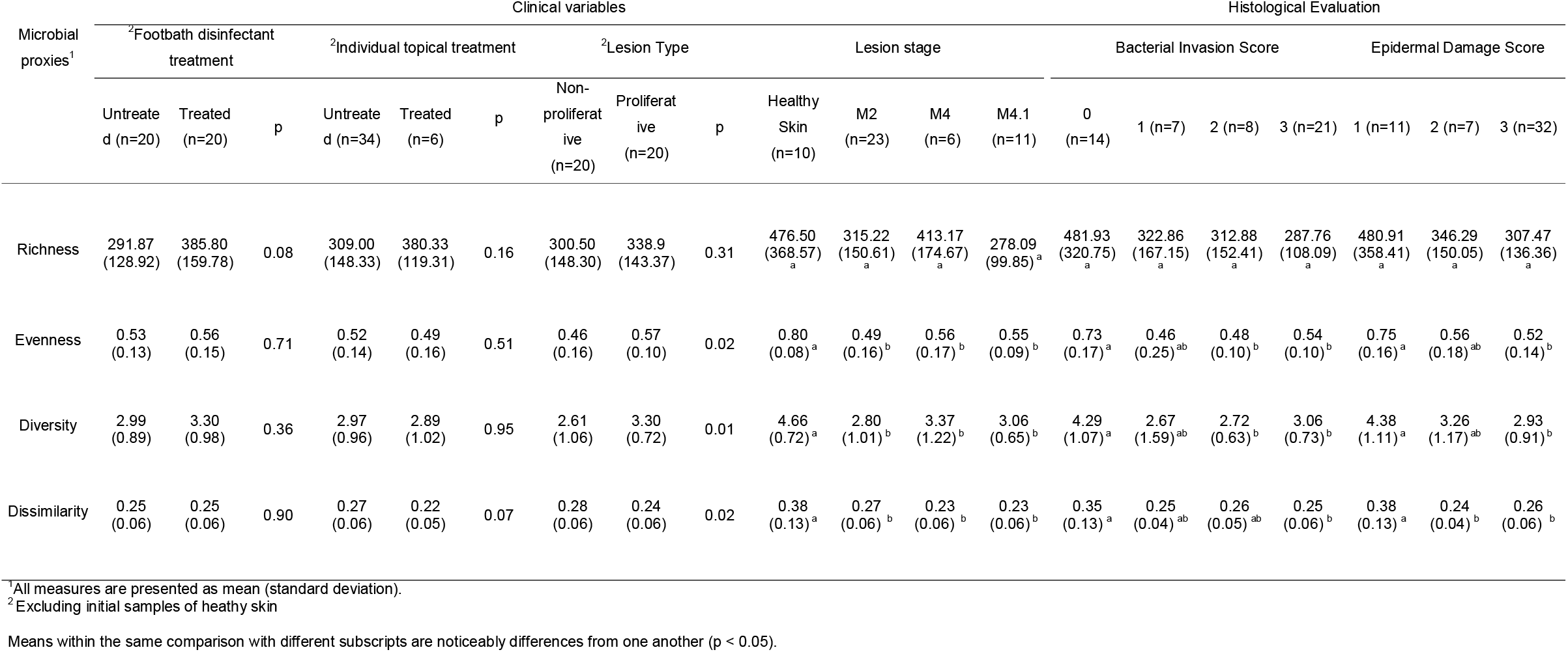
Effect of different clinical variables on the microbial proxies studied in 10 dairy cows with bovine digital dermatitis lesions followed over 45 days.

Regarding the composition of the microbiota and in terms of relative abundance, the most frequent phylum found among the healthy skin samples were Firmicutes (49%), Actinobacteria (15%), Bacteroidetes (13%), Spirochaetes (11%), and Proteobacteria (5%). Otherwise, the most frequent phylum found in DD lesions at day 0 were Spirochaetes (54%), Bacteroidetes (25%), Firmicutes (14%), and Tenericutes (3%) (Figure S1).

A visual description of the structure of DD lesion microbiota over time is provided in Figure S2. The microbiota from healthy skin (day 0) and DD lesion at day 45 showed a more dispersed pattern compared to the other time points (Figure S2 A). Nevertheless, when comparing all-time points together, no major visual differences or cluster formation were evidenced (Figure S2 B).

Figure 2A shows the microbiota according to whether or not the footbath disinfectant was used during the study period (DD samples from days 15, 30 and 45). No distinctive clustering is observed. Regarding the use of individual topical treatment, the treated samples seem to cluster separately with an obvious reduction of Spirochaetes and, in particular, Fusobacteria were placed completely outside the clusters (figure 2B). The scatter plot of the lesion type suggests a distinctive cluster for proliferative and non-proliferative lesions (Figure 2C). Finally, based on the clinical score, only the microbiota of healthy skin appears to separate from the cluster formed by DD lesions. Nevertheless, each stage of the lesion seems to draw distinctive scatter plots (Figure 2D). Otherwise, according to the histological evaluation, scores 0 and 1 seems to cluster separately from scores 2 and 3 when comparing bacterial invasion scores (Figure S3 A). Likewise, score 1 and 2 seems to cluster separately from score 3 when comparing epidermal damage scores (Figure S3 B).

**Figure 2.**
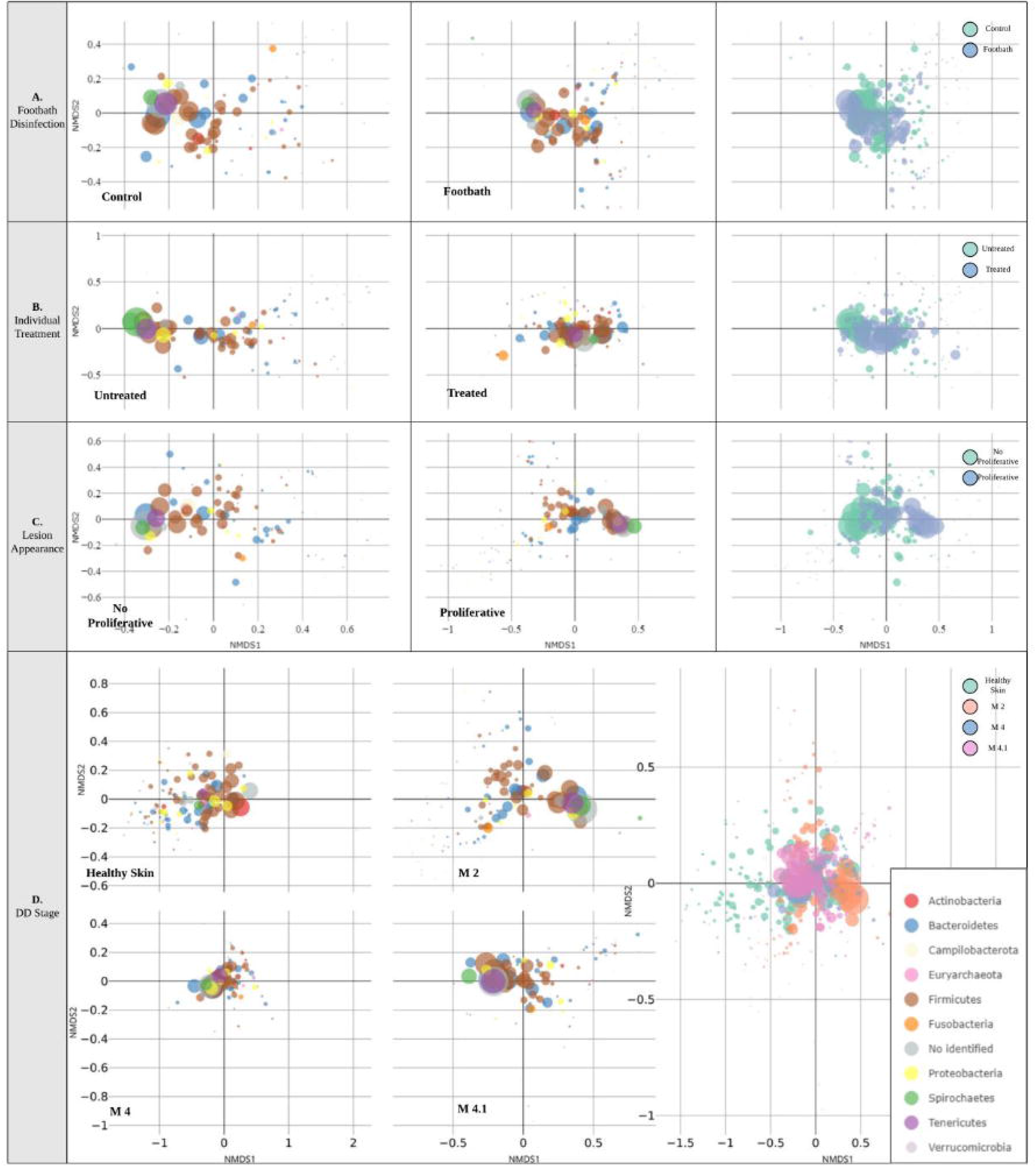
Nonmetric multidimensional scaling (NMDS) scatterplot of Bray-Curtis dissimilarities of the skin microbiotas from feet suffering bovine digital dermatitis (DD) lesions. Microbiotas are visualized by dots representing the amplicon sequence variants (ASV) present in the samples according to the usage of the footbath disinfectant (A), the usage of individual topical antibiotic treatments (B), the lesion type (C) and the M5 score lesion score (D). The size of the dots represents the mean relative ASV abundance and are colored according to their taxonomic classification at the Phylum level.

Finally, the dynamics of DD microbiota composition were explored through a differential abundance analysis (Figure 3). For the baseline comparison between healthy skin and DD lesions at day 0, 40 ASVs significantly reduced their abundance and 58 increased their abundance. Indeed, compared to healthy skin, the microbiota of DD lesions at day 0 have increased abundance of *Treponema* spp. (∼4 to ∼28 FC), *Fretibacterium* spp. (∼5 FC), *Mycoplasmopsis* spp. (∼6 FC) and *Proteus* spp. (∼21 FC). Decreased abundances of *Campylobacter* spp. (∼17 FC), other unidentified Spirochaetes (∼17 FC), and different genera of Proteobacteria (∼4 to ∼18 FC) and Tenericutes (∼5 FC). Changes in both directions were detected on different genera and other unidentified Bacteroidetes (∼-20 to ∼10 FC) and Firmicutes (∼-22 to ∼22 FC) abundances. When comparing the abundances of DD samples from day 15 to day 0, the abundance of only 18 ASV changed significantly. In detail, there was a reduction in *Treponema* spp. (∼16 FC) and *Proteus* spp. (∼21 FC), and an increase in Actinobacteria (∼16 FC). Different genera and other unidentified Bacteriodetes (∼ -5 to ∼18 FC) and Firmicutes (∼ -28 to ∼17 FC) change in both directions. For the comparison between day 30 and day 15, the abundance of 34 ASV changed significantly. We noticed a reduction in *Oligella* spp. (∼19 FC) and an increase in *Campylobacter* spp. (∼4 to ∼26 FC) and other unidentified Spirochaetes (∼34 FC), Proteobacteria (∼24 FC) and ASV (∼40 FC). In particular, changes in both directions were observed in *Treponema* spp. (∼-25 to ∼24 FC) and for different genera and other unidentified Bacteriodetes (∼ -24 to ∼33 FC) and Firmicutes (∼-20 to ∼34 FC). Finally, the abundance of 22 ASV changed significantly when comparing form day 45 to day 30 microbiota. We noticed a reduction in *Treponema* spp. (∼6 to ∼25 FC) and an increase in *Oligella* spp. (∼21 FC) and other unidentified Proteobacteria (∼7 FC). Changes in both directions were observed for different genera and other unidentified Bacteriodetes (∼-10 to ∼26 FC) and Firmicutes (∼ -34 to ∼24 FC).

**Figure 3.**
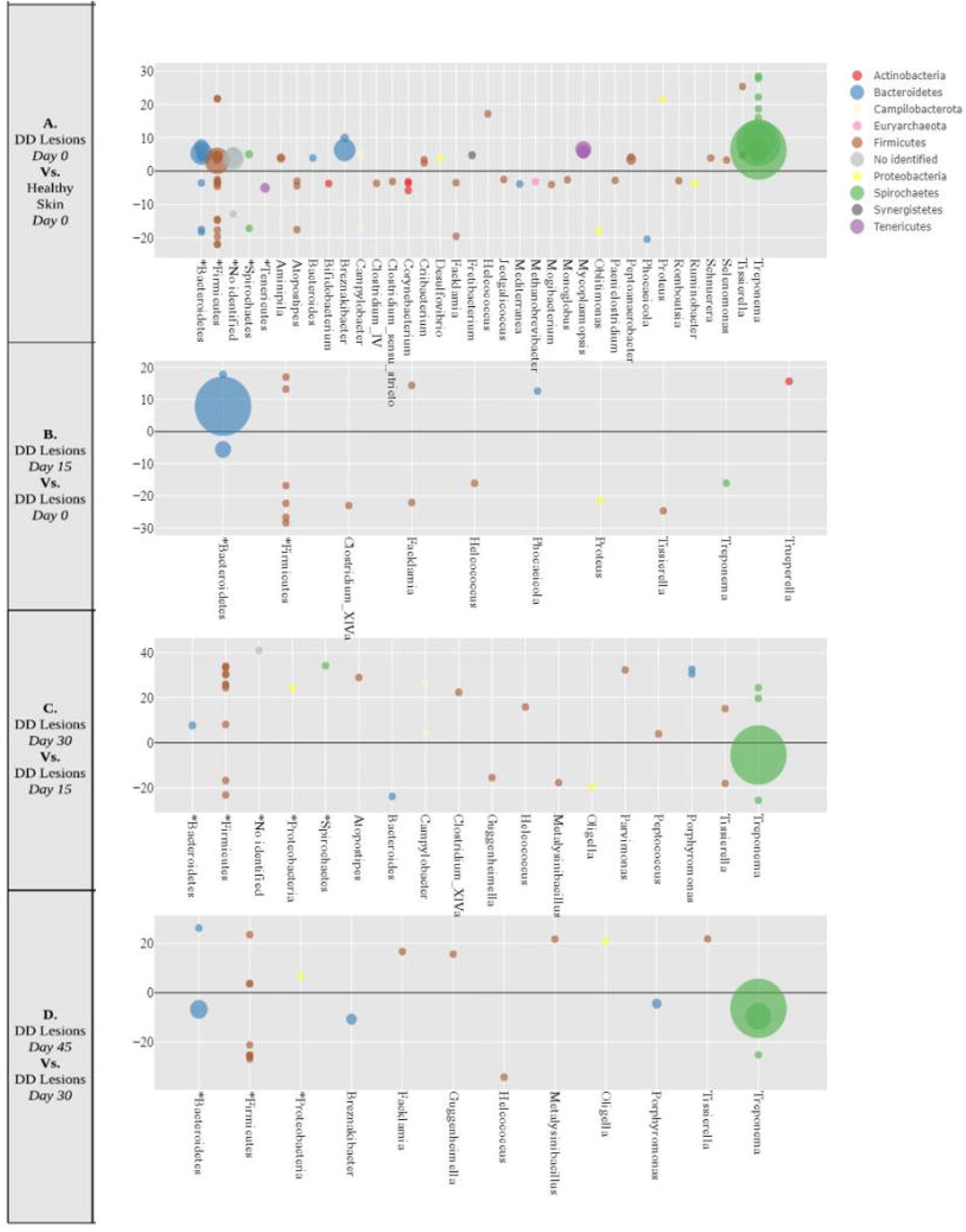
Differential abundance of amplicon sequence variants (ASV) present in skin microbiotas from feet suffering bovine digital dermatitis (DD) lesions. (**A**) DD lesions versus healthy skins at day 0, (**B**) DD lesions at day 15 versus DD lesions at day 0, (**C)** DD lesions at day 30 versus DD lesions at day 15, (**D**) DD lesions at day 45 versus DD lesions at day 30. Bubbles represent the ASV classified at the genus level (x-axis) that showed noticeably changes (*p value* < 0.1) on a log2-scale (y-axis). The size of the bubbles represents the mean relative ASV abundance and are colored according to their phylum-level classification. Unclassified ASV at the genus level are gruoped at the Phylum level (*).

## Discussion

The present investigation describes the differences between microbiotas from healthy skin and those affected by DD lesions under field conditions. Additionally, for the first time, the microbiota dynamics of DD lesions were investigated during a period of 45 days exploring the putative impact of recognized factors, such as the use of individual and collective treatments. As expected, DD lesions microbiota were dominated by *Treponema* spp. and their configuration was largely different from healthy skin in terms of diversity, structure, and composition. The DD lesions microbiota also evidenced significant changes in its structure, in particular the populations of *Treponema* spp., *Oligella* spp., Spirochaetes, Bacteriodetes and Firmicutes constantly changed over time. Although the footbath disinfectant and the individual topical treatment used in this study have recognized DD healing potential based on their antimicrobial properties, no major impact on microbiota characteristics was associated with their use over time. Other important determinants, such as foot hygiene or the farm environment, may have altered the effectiveness of these strategies. In addition, we detect significant differences between non-proliferative and proliferative lesions microbiotas. We recorded a partial recovery of the microbiota diversity trough the 45 days of follow-up. Our findings reveal important dynamic changes on the abundances of *Treponema* spp. during the study period. For the first time, this investigation provided insight into how the microbiota of DD lesions may change over time under field conditions.

Our findings support the polymicrobial nature of DD highlighted in previous investigations using sequencing and culture–based approaches. It is in line with findings from previous research signaling *Treponema* spp. as the main microorganism related to DD lesions(5, 8, 9). As expected, the microbiotas gathered in this study share a similar structure with microbiotas previously described on healthy skin and DD lesions from dairy cattle, respectively(5, 20). In detail, the main bacteria linked to DD lesions in previous investigations that were again identified by our findings include: *Treponema* spp., *Acholeplasma* spp.(5, 8, 9), *Porphyromonas* spp., *Prevotella* spp.(12), *Corynebacterium* spp.(9), *Tissierella* spp., *Mycoplasmopsis* spp.(8, 21) and some unclassified Bacteriodetes, Firmicutes and Proteobacteria(11, 12, 20). On the contrary, some bacteria previously associated with DD, such as *Dichelobacter nodosus*(14, 22) was not linked to lesions in this study. Although *Fusobacterium* spp. have been previously related to DD lesions(5, 8, 9), our findings indicate that its presence seems to be related to a particular farm environment rather than to the disease microbiota dynamics. The present study also found an over-representation of *Fretibacterium* spp. in DD lesions compared to healthy skin, an anaerobic bacterium not previously implicated DD but, like different *Treponema* spp, related to human periodontitis(23). Our findings coincide with the results of a recent meta-analysis revealing the main role of *Treponema* spp., *Mycoplasmopsis* spp. on DD pathogenesis(24). Indeed, we recorded significant changes in the abundances of a large number and diversity of genera in DD lesions compared to healthy skin. Nevertheless, future studies are needed to elucidate which specific genera might be related to disease occurrence, regional particularities of the sampled farms, or sampling contamination. Similarly, we recorded consistent changes in the abundance of *Oligella* spp. over time, a bacterium previously reported in ruminant milk microbiota(25) but not yet associated with DD and which might be an active part of the farm environment. However, as these are the first skin microbiota assembled from dairy cattle in France, no regional comparison can be performed. Additionally, the fact that previous studies have not linked this bacteria genus to DD lesions might be explained by the original statistical procedures implemented in the analyses of our present study. Indeed, the relative abundances of every sample were calculated after the standardization of the observed taxonomic groups according to their variance and not through procedures implemented previously such as the rough normalization which may entail the loss of valuable information(26). More in detail, when using rough normalization procedures for data analyses, information from samples with a scarce number of sequences is excluded from the dataset. In specific cases, such as healthy skin samples, the exclusion of this information may conduct to neglect the healthy nature of the sample which could potentially have low numbers of sequences in comparison to diseased samples. The within-cow design of this study offered the possibility of exploring the dynamics of microbiotas according to the clinical evolution of each animal. Indeed, the simple comparison between microbiotas from DD lesions obtained from different animals at diverse states may underestimate or neglect the impact of individual and environmental factors (i.e. foot conformation or barn hygiene) on the clinical outcome of lesions and their microbial configuration.

One of the main limitations of this study was related to the difficulties encountered in interpreting microbiotas obtained from an anatomical region which inevitably is in close contact with the ground and thereby with dirtiness. Indeed, the distinction of pathogens among environmental microorganisms is a challenge. Furthermore, due to the nature of our molecular approach, our study not provide evidence of bacterial viability. We have tried to reduce this limitation by the meticulous washing and scrubbing procedures performed during the biopsy collection, which in turn may have altered the skin microbiota. However, colonization of tissues by opportunistic pathogens could be favored by the successive incisional biopsies performed during the follow-up of the lesions. Moreover, depending on the lesion size successive biopsies in a same location could restraint the expected healing of the skin. Another limitation of the present investigation was related to the small sample size studied. Larger samples may highlight with more precision the benefit of control strategies for DD in increasing the microbiota diversity of the skin. As several risk factors at the farm level are involved with the disease, such as hygienic conditions, differences in the microbial profile according to the farms were expected. However, the small number of farms (n=5) and the small number of individuals sampled within farm restrain the exploration of factors related to farm hygiene. In this study, the changes in the microbiota allowed a clear distinction between the diseased (M2-4-4.1) and healthy (M0) states, but the potential impact of individual or collective treatments on the microbiota was not detected. In contrast to previous studies(27), for the present investigation the potential utility of NGS tools to measure the potential effectiveness of control strategies seemed inferior or at least different from standard observation methodologies such as DD stages or lameness scoring. In terms of ethics, feasibility, and more importantly precision, invasive methodologies (biopsies) are limited in large samples and, as pointed above, entails a challenge for the interpretation. Another point concerns the fact that the effect of both, the footbath disinfectant and the individual topical treatment, could be highly affected by the exacerbated anti-inflammatory response of the animals after each biopsy. Therefore, measuring local immune responses may enhance the precision of the true effect of control interventions. Regarding the histological evaluation of the samples, this additional assessment clearly allowed the distinction of the bacterial invasion advance between the samples. Although bacterial invasion (BIS) was in concordance with the clinical evaluation of the samples and the dynamic evolution of the microbiota abundances, the epidermal damage (EDS) was more difficult to interpret due to the potential effect of the recurrent biopsy procedures.

Among the different clinical factors studied, the structure and diversity of microbiotas from DD lesions were not affected by the usage of individual topical treatment. This could be explained by the short and not persistent effect of this strategy over time(28). Otherwise, the findings of this study indicate noticeable differences between the microbiotas of non-proliferative and proliferative DD lesions. The presence of proliferative tissues is associated with chronical processes and thereby to lesions prone to relapse(29). The hyper-proliferation of tissues surrounding the lesion may alter the environment of the skin and the topical activity of footbath and individual topical treatment, thus influencing a distinctive microbiota(30–32). Recently, *Treponemas* spp. have also been involved in some non-infectious diseases inducing lameness(33). Further studies are needed to evaluate the role of microbiota in the dynamics of those non-infectious lesions that are related to the same risk factors recognized for DD. Other recognized factors influencing the occurrence of DD lesions at the individual level, such as parity or days in milk, were not studied due to the small number of cows evaluated at every time point and the large variability among them.

Beyond the recognition of the pathogenic role of some specific bacteria, exploring the microbiotas from DD lesions at the community level may highlight how the dynamics of skin microbiota determine the persistence and the occurrence of DD lesions. Indeed, the 16*S* rRNA gene analyses may capture broad shifts in community diversity over time, but with limited resolution and lower sensitivity compared to metagenomic data. Nevertheless, the results of this investigation reveal the dynamic nature of microbiotas from DD lesions and highlight the potential importance of specific bacteria in the progress of the lesions over time. Future studies could benefit from shotgun sequencing tools, improving sensitivity in recognizing pathogens and inhabitants and defining their specific role in disease. Therefore, further investigations linking the skin microbiota to different herd management practices might enhance the current understanding of the disease and promote the development of successful control strategies.

In conclusion, we have explored the dynamics of the skin microbiota in feet affected with DD lesions. The diversity and structure of microbiotas from DD lesions were different from those of healthy skin but did not vary according to the usage of the footbath disinfectant or the individual topical treatment. The microbiota of proliferative lesions displayed a different structure and diversity compared to non-proliferative lesions. Concerning the dynamics of DD lesion progression, we confirmed the putative role of *Treponema* spp. and open new questions about the potential role of *Mycoplasmopsis* spp., *Oligella* spp. and other Spirochaetes, Bacteriodetes and Firmicutes. Further studies are needed to understand how risk factors associated with DD at the farm and individual level may affect the interactions between the bacteria comprising the skin microbiota, and how these interactions may determine the clinical evolution of DD lesions.

## Methods

### Ethics statement

The research protocol was reviewed and approved by the Ethics Veterinary Committee in Clinical Research and Epidemiology from the Veterinary School of Nantes, ONIRIS, France (CERVO, France) (registered number: CERVO-2016-12-V).

### Study population

Five dairy farms participating simultaneously in a controlled clinical trial were included in the study(34). The herds were composed exclusively of Holstein cows to limit the putative breed effect. The herds were housed in cubicles and on average 90 cows were milked twice a day in a rotatory or conventional milking-parlor. The prevalence of active DD lesions at the start of the study ranged from 20% to 40%.

During the study period (45 days), farmers were allowed to detect and then treat severe cases of DD by using 2 applications of oxytetracycline (30 mg/ml) (Oxytetrin™, MSD Animal Health/Intervet, France) 2 days apart. For the current investigation, from each farm, the owners have chosen two animals suffering from a rear DD lesion. Therefore, the study population consisted of 10 different cows affected by DD. For each cow, a single lesion was recorded and follow-up during the study period. Five of the cows did not receive any collective disinfectant treatment on the foot suffering the DD lesion recorded and the remaining 5 feet received a collective disinfectant treatment through a footbath (Pink-Step™, Qalian). The frequency of administration of the footbath was of 2 days (4 consecutive milkings) every week of the study period. The cows included in the study were between their first and fifth parity, and their days in milk ranged from 3 to 378 at the start of the study.

### Follow-Up and Data Collection

Farms were visited by 2 investigators trained by practical lessons to practice the biopsies and score the lesions. All lesions sampled were photographed and scored using the M5 score system(32). At every sample time, we recovered information about the eventual administration of any concomitant treatment between the samplings. To perform the biopsies every cow was carefully restrained in a trimming chute. The skin area was washed with water and then local anesthesia was provided using Procaine 2% (Procamidor™, Axience, France). Thereafter, the area was rinsed and brushed with a PBS solution before performing the sample with a sterile biopsy punch (6 mm). The incisional samples were again washed with sterile PBS solution, cut tangentially into two approximately equal parts. One half was stored in 10% neutral buffered formalin solution for histological evaluation, and the other half was stored in sterile 2.0 ml microcentrifuge tubes at -20°C for DNA extraction and sequencing analysis. Finally, an aerosol bandage of aluminum (Aluspray™, *Vetoquinol*, France) was sprayed directly over the incisional biopsy wound of each cow. Additionally, in the case of pain detected by farmers, NSAIDs were provided after the biopsy sampling.

For every cow, five skin biopsies of the affected foot were performed at 4 different times. Only for the first sampling, two biopsies were taken at the same foot, one sample on the healthy skin, and the second on the DD lesion. The biopsy sampling start immediately before the application of first footbath. Thereafter, 3 subsequent biopsies of DD lesions were taken every 15 days. This time interval was chosen to allow the partial healing of the skin. The location of each biopsy lesion intended to be taken as close as possible to the previous sampling sites. In total 50 biopsies were recovered.

### Histopathology

The paraffin sections of every tissue sample were stained with *Hematoxylin-Eosin-Safran* (HES) for the general histopathological evaluation and with *Warthin Starry Silver (WSS)* to evaluate the presence and degree of spirochaetal invasion. The degree of epidermal damage (EDS) was assessed according to a modified score system described in a previous study(14). A score of 1 was defined as normal skin or presenting mild acanthosis and hyperkeratosis without any other significant microscopic lesions associated and without bacteria. Score 2 was defined as skin with marked acanthosis and hyperkeratosis, without erosion or ulceration, with or without pustules, with or without hemorrhages and erythrocytic crusts with or without dermal fibrosis. Finally, score 3 was defined as skin presenting a segmental epidermis, localized to extensive, necrotizing to necrosuppurative epidermis with balloon degeneration of epithelial cells, necrotic vasculitis and intralesional bacteria (spirochaetes, bacilli, and coccobacillus). Otherwise, the total bacterial invasion score (BIS) encompassed also the spirochaetes identified by the WWS staining. The score 0 represented the absence of invasive bacteria; score 1, a low number of invasive bacteria; score 2, a moderate number of invasive bacteria; score 3, a high number of invasive bacteria.

### DNA Extraction, 16S rRNA amplification, illumina MiSeq sequencing

Bacterial DNA extraction was enhanced adding to every sample 20 μl of proteinase K, 180 μl of tissue lysis buffer, and 40 μl of lysozyme (Qiagen, Valencia, CA, USA). Thereafter, all the samples were incubated for 12 h at 56°C and then processed directly for DNA extraction using the Powerlyzer Powersoil Kit (Quiagen, Valencia, CA) according to the manufacturer’s instructions. Extractions were performed in rounds of 10 samples and in every round, an additional empty tube was processed in parallel to serve as a negative extraction control. Finally, the resulting supernatant of each sample was transferred to a labeled microcentrifuge tube and stored at -20°C The Quant-iT dsDNA Assay Kit (Thermo Fisher Scientific, MA, USA) was used to measure DNA concentrations prior to PCR amplification. The V4 region of the bacterial 16S rRNA gene was amplified by PCR, using Illumina Nextera XTIndex Primer 1 (N7xx) and Primer 2 (S5xx), where barcodes were unique to each sample. The PCR protocol consisted of an initial denaturation at 95 □C for 10 minutes followed by 40 cycles of 95 □C for 30 seconds and 60 □C for 1 minute and the real-time fluorescence data acquisition occurred at the end of each annealing/extension phase. All PCR assays use the same PCR cycling conditions allowing parallel testing of all PCR assays. The amplicons were extracted and purified from the gel using the Zymo gel extraction kit according to the manufacturer’s instructions and quantified using the Quant-iT dsDNA Assay Kit (Thermo Fisher Scientific, MA, USA). Finally, the purified amplicons were sequenced using the MiSeq reagent kit v2 (13 cycles) on the MiSeq platform (Illumina, Inc., CA, USA).

### Sequencing data processing

The generated 16S rRNA raw gene sequences were preprocessed to remove any remaining, barcodes or adapters using Trimmomatic(35). Raw reads were then processed using R(36), implementing the established DADA2 workflow(37). The amplicon sequence variants (ASV) in each sample were inferred using the “divisive amplicon denoising algorithm”, a parameterized model of substitution errors to distinguish sequencing errors from real biological variation(38,39). The inferred forward and reverse sequences were then merged, removing sequences that did not perfectly overlap and subsequently chimeras were removed. To each ASV a taxonomic label was assigned through a Bayesian classifier using the Ribosomal Database Project (RDP) v18 training set(40, 41). The phylogenetic relatedness of the ASV identified was calculated using the R package *decipher* (42) and, spurious ASV not occurring in two or fewer samples were removed. Finally, the ASV were agglomerated according to a cophenetic distance inferior to 0.03. Sequencing data have been deposited in the MG-RAST (https://www.mg-rast.org) database and are publically available at: https://www.mg-rast.org/linkin.cgi?project=mgp80659.

### Statistical analyses

The statistical analyses were performed using R (36). The analysis strategy consisted of the following 4 steps: i) Describe the dynamics of the skin microbiota on feet with DD lesions using different microbial proxies of diversity and community structure, ii) explore whether the studied microbiotas were affected by different factors that have been previously identified as impacting the clinical course of DD lesions (clinical variables), and iii) identify differential ASV that may affect the course of DD lesions over time.

The clinical variables explored were the usage of the footbath disinfectant, the usage or not of individual topical antibiotic treatments, the M5 stages recorded, the lesion type (proliferative or not), and the histopathologic evaluation (degree of bacterial invasion and lesion scoring). The microbiotas were charcaterized using four different microbial proxies including: richness (count of observed ASV), evenness (Pielou index), diversity (Shannon index), and dissimilarity (43). More in detail, while the Pielou index measures the evenness of ASV abundances and how abundances are distributed, the Shannon index determines the microbial diversity controlling by the richness (44). The dissimilarity was explored by calculating the Bray-Curtis distances according to the different clinical variables studied. All microbial proxies were compared between the clinical variables studied using Kruskal-Wallis and pairwise Mann–Whitney U tests. The impact of the clinical variables on the structure of microbiotas over time was visualized using the Bray-Curtis distances and nonmetric multidimensional scaling ordination (NMDS). In the scatter plots created, high dissimilarity of microbiotas was represented by long distances between dots, whereas microbiotas with a similar composition were clustered together.

Finally, the dynamic on DD microbiota composition was explored by identifying specific ASV changing over time through a differential abundance analysis using the R package *DEseq2* (45). *DEseq2* uses a negative binomial distribution and implement the variance stabilization transformations of the ASV read counts identified in the raw data set to account for overdispersion. A single model was fitted including the main effects of the sample time (day 0, 15, 30 and 45) and independent hypothesis weighting (IHW)(46) was used to corrected for multiple comparisons such as the dependencies between the multiple samples within the same cow. Therefore, significant log 2 fold change (FC) (p-value ≤ 0.1) on the ASV counts were explored between the microbiotas of: i) healthy skin Vs. of DD lesions at day 0, ii) DD lesions at day 0 Vs. DD lesions after 15 days, iii) DD lesions at day 15 Vs. DD lesions after 30 days and, iv) DD lesions at day 30 Vs. DD lesions after 45 days. All figures were created with the R packages *ggplot2* and *plotly* (47, 48).

## Acknowledgments

The authors acknowledge the farmers and veterinarians who have contributed to this study. Additionally, we thank G. Puel (Oniris, Nantes, France) for helpful participation as investigator, for his time and motivation. This study was supported as part of a Ph.D. studentship by Qalian (Neovia group, Segré, France) and the “Association Nationale de la Recherche et de la Technologie” (ANRT, Paris, France).

## Author contributions

J.M.A., D.D., K.O., A.R., N.B., and R.G. designed the study. J.M.A., S.L., D.D., and K.A. analyzed the data. J.M.A. and R.G. interpreted results. J.M.A. and R.G. drafted the manuscript. All authors revised critically the manuscript for important intellectual content and approved the final version to be published. The authors acknowledge the farmers and veterinarians who have contributed to this study.

## Figure Legends

**Figure S1**. Relative abundances of the ASV present in the samples according to the sampling day. Taxonomic classification at the phylum level (A) and at the genus level (B).

**Figure S2**. The composition of skin microbiotas from feet suffering bovine digital dermatitis (DD) before and across the 45 days of study follow-up. Microbiotas are visualized by dots representing the amplicon sequence variants (ASV) identified in the samples. The size of the dots represents the mean relative ASV abundance and are colored according to their taxonomic classification at the Phylum level. The ordination was constructed using a Bray-Curtis matrix and Nonmetric multidimensional scaling (NMDS).

**Figure S3**. Nonmetric multidimensional scaling (NMDS) scatterplot of Bray-Curtis dissimilarities of the skin microbiotas from feet suffering bovine digital dermatitis (DD) lesions. Microbiotas are visualized by dots representing the amplicon sequence variants (ASV) present in the samples according to the histological evaluation for bacterial invasion (A) and epidermal damage (B). The size of the dots represents the mean relative ASV abundance and are colored according to their taxonomic classification at the Phylum level.

