## Supplemental Figure 1-3 for "Skin microbiota dynamics of dairy cows affected by digital dermatitis"

**Figure S1.** Relative abundances of the ASV present in the samples according to the sampling day. Taxonomic classification at the phylum level (A) and at the genus level (B).

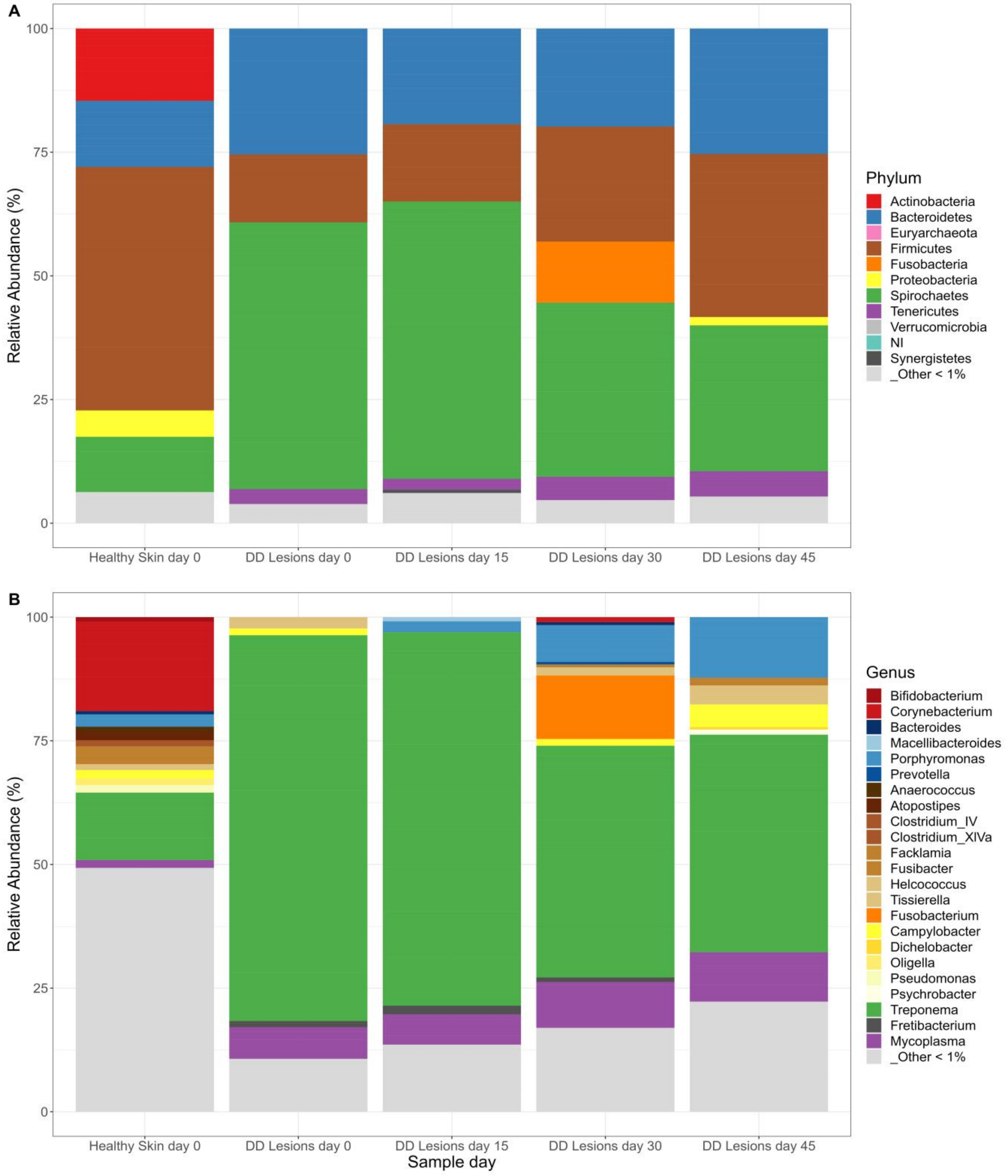

**Figure S2.** The composition of skin microbiotas from feet suffering bovine digital dermatitis (DD) before and across the 45 days of study follow-up. Microbiotas are visualized by dots representing the amplicon sequence variants (ASV) identified in the samples. The size of the dots represents the mean relative ASV abundance and are colored according to their taxonomic classification at the Phylum level. The ordination was constructed using a Bray-Curtis matrix and Nonmetric multidimensional scaling (NMDS).

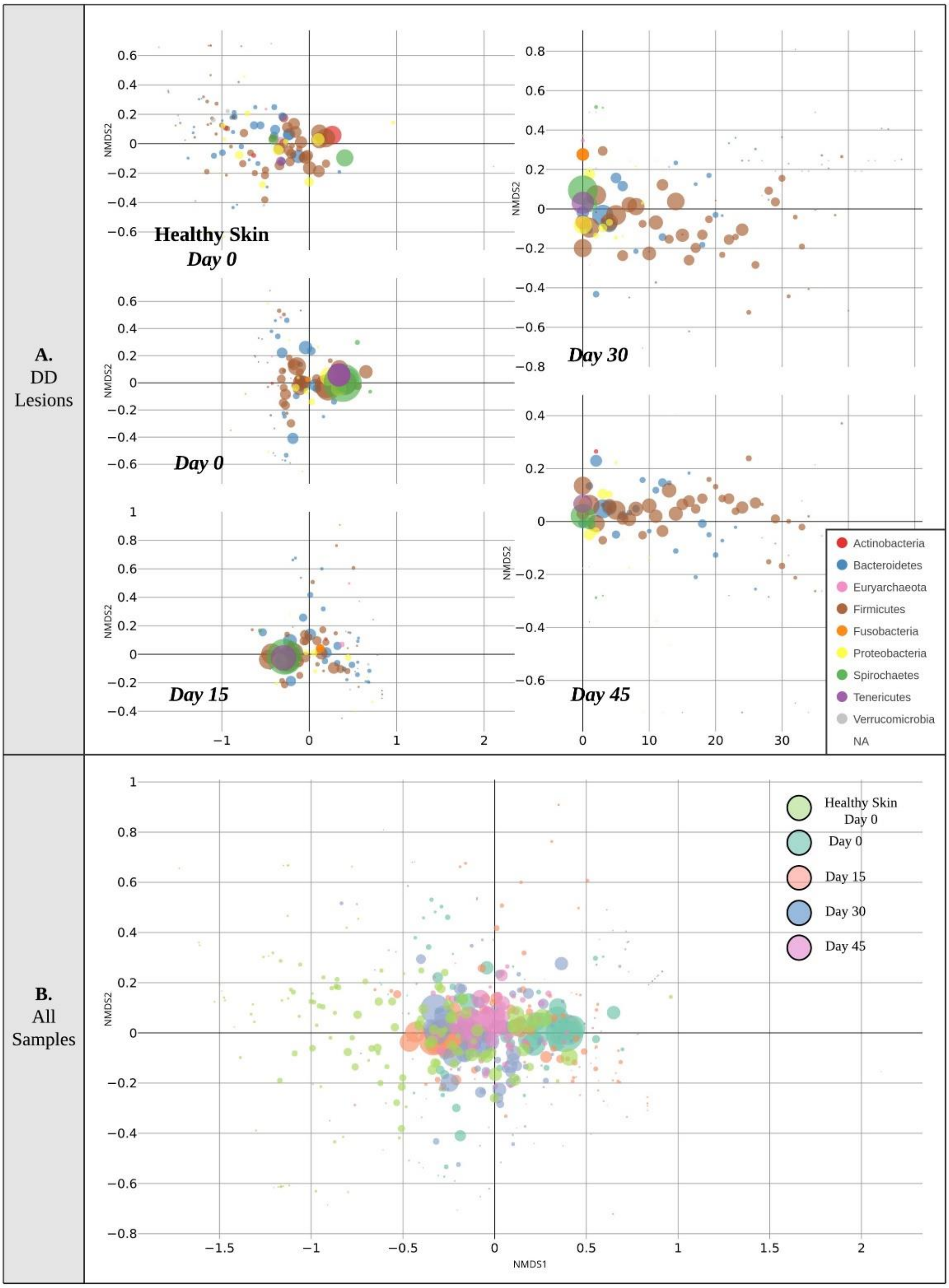

**Figure S3.** Nonmetric multidimensional scaling (NMDS) scatterplot of Bray-Curtis dissimilarities of the skin microbiotas from feet suffering bovine digital dermatitis (DD) lesions. Microbiotas are visualized by dots representing the amplicon sequence variants (ASV) present in the samples according to the histological evaluation for bacterial invasion (A) and epidermal damage (B). The size of the dots represents the mean relative ASV abundance and are colored according to their taxonomic classification at the Phylum level.

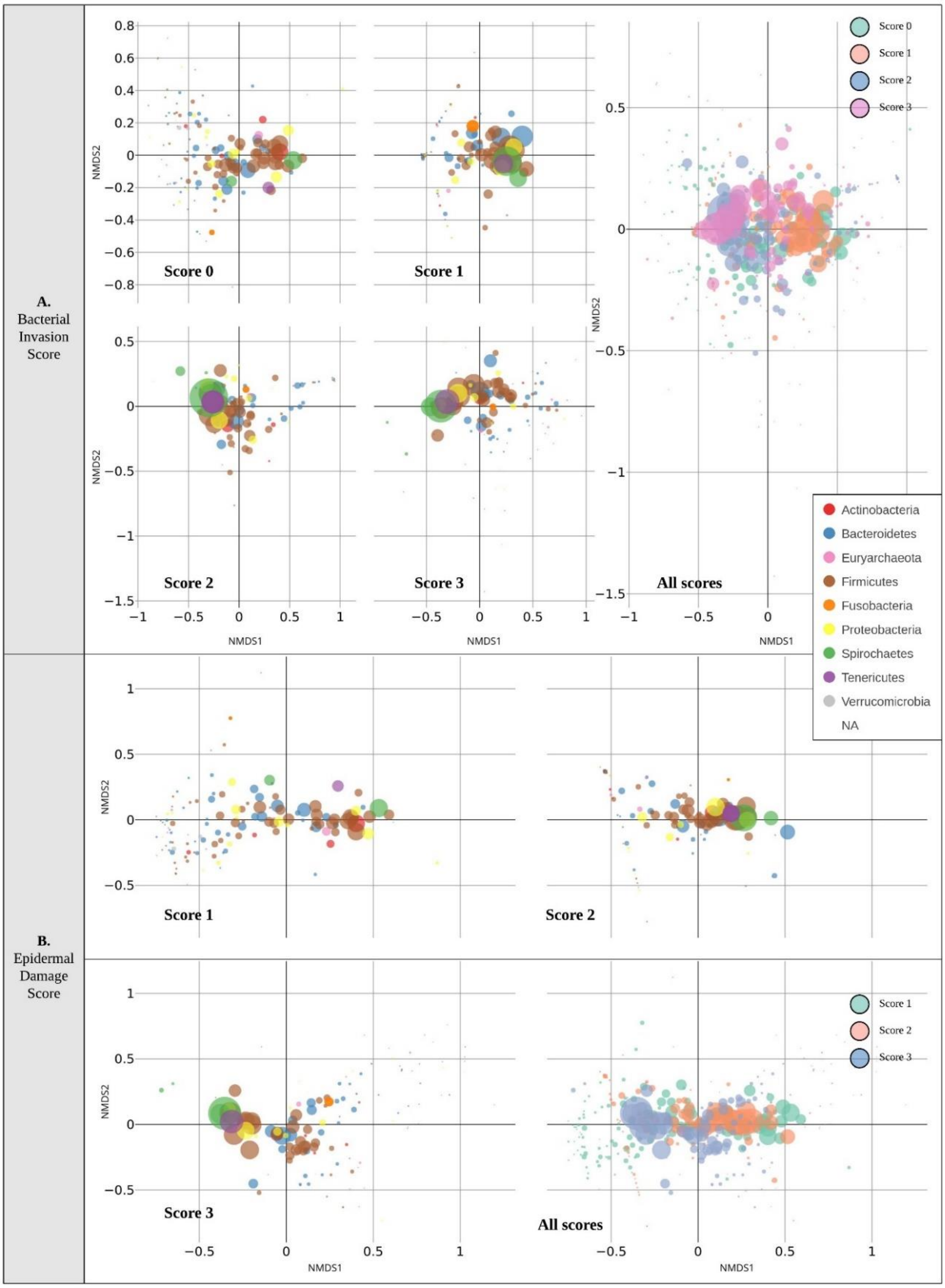
